# Graph Neural Networks (GNNs) for Protein-Ligand Interaction Prediction

**DOI:** 10.64898/2026.04.23.720519

**Authors:** Subhasankar Khilar, Elamathi Natarajan

## Abstract

Predicting protein-ligand interactions in the modern drug discovery has revolved from the involvement of artificial intelligence and structural bioinformatics using Graph Neural Networks (GNNs). The limited explainability of GNN models presents an important encumbrance in biomedical research, but it has achieved a high degree of accuracy in determining and identifying binding affinity and active compounds, as evidenced by [1] [2] [3] [4]. Here this research focuses on the interpretation of protein-ligand interactions at a molecular level, a rapidly developing area within Graph Neural Networks (GNNs). Now days modern study handling techniques such as visualization techniques, attention mechanism and model-based feature ascription by model to boost, and make robust and decrease false predictions on binding. Along with some approaches include like graph pooling strategies, message-passing optimization, self-supervised learning, transfer learning and contrastive learning are rapidly utilized to enhance the representative learnings. Furthermore, integration of molecular docking simulations, hybrid deep learning architectures and protein language model gives more reliable & biological predictions of protein-ligand interactions.

That focuses on given process that identifies key ligand atoms and binding residues, as well as physicochemical factors influencing affinity, through chemical thought processes. Here this research work identified the challenges of developing biologically significant explanations, transparency, and the corollary dataset biases on interpretability. The research work conducted an in-depth investigation into the consolidation of protein language models to establish more reliable pathways for future research, examining hybrid architectures, transparent and energy-efficient GNNs, and scientifically grounded AI models for drug discovery. My research work highlights that XGNNs establishes a connection between Deep Learning and Biochemical expertise with increased confidence, which will enhance the accuracy of predictive models and computational models.

## I. Introduction

### 1.1 Background of Drug Discovery

The method of drug discovery is a highly complex, expensive and time-consuming process. Along with it needs identification of potential drug targets that can interact with target proteins. The conventional drug discovery methods like in-vivo and in-vitro studies they need a fixed number of resources and responsible for high rate of failure. [5] [6]

On other hand a crucial step in this is, understanding the Protein–Ligand Interactions (PLIs), where a drug molecule or ligand binds to a target protein to produce a biological response. The discovery of potential drug candidates and devaluation of experimental costs can be done through accurate prediction of these interactions.

In past, the computational techniques such as molecular docking and computer-aided drug design (CADD) were developed in the 1980s to accelerate these techniques allow for simulation of interactions between molecules. Although these techniques have some limitations in understanding the complex molecular structures and interactions. [7]

### 1.2 Role of Artificial Intelligence in Bioinformatics

With the advent of Artificial Intelligence (AI), especially Machine Learning (ML) and Deep Learning (DL), there has been a tremendous change in the field of Bioinformatics. This is possible only because of AI, which is capable for analysing a large-scale of biological information and making predictions about the molecules. [4]

The recent advancement in this field is focused on learning representations directly from structures. This has enhanced the predictions of protein-ligand binding affinity. These techniques reduce computational cost and make predictions more efficient.

AI-based technique provides different type of tools for interpreting, interpreting and extracting meaningful insights from huge and high-dimensional biological data. Machine Learning (ML) techniques gives predictive modelling by analysing from existing data, whereas Deep Learning (DL) techniques further increases its capability by automatically learning features from data without requiring human intervention. This allowed to researchers to move from descriptive models to predictive models in biological research. [8]

In the field of drug discovery, AI played a crucial role to play in different stages of drug discovery process. Which includes target identification, virtual screening, toxicity prediction and lead optimization etc. However, in the drug discovery domain the most momentous contribution of AI has been in the prediction of protein-ligand interactions, which are mostly essential in understanding the efficacy of the drug. However, the conventional computational drug discovery approaches, including molecular docking and molecular simulation, along with involve a high computational cost and may not be sufficient to explain the interactions between the drug and the target. [9]

Moreover, the AI-based approaches provide a cost effect in the drug discovery domain that allows the high-throughput screening of the large number of compounds, thereby saving a lot of computational cost and time that would be spent in the experimental analysis of the large number of compounds.

### 1.3 Introduction to Graph Neural Networks (GNNs)

Graph Neural Networks (GNNs) have been developed as a promising and advanced deep learning technique particularly designed for the handling of non-Euclidean data structures in which relationships between entities are not linear and cannot be effectively handled by conventional grid-based structures such as images or sequences. In some real-world biological systems, the data tends to exist in a non-linear format as networks rather than in linear or grid-based formats, making GNNs particularly attractive for use in bioinformatics. [10]

In the context of molecular biology and drug discovery, both ligands & proteins can be effectively modelled as graphs. A molecule is only not a set of atoms; it is a system in which, relationships between atoms determine the molecule’s chemical and biological properties. In the context of the proposed GNNs approach:

- Nodes in the graph represent atoms in the ligand, whereas amino acid residues in the protein are represented by nodes.
- Edges in the graph represent the bonds between atoms.

Unlike traditional neural networks that process information independently, whereas GNNs process information by passing information across the nodes through their edges. Another thing is, this allows a node to gather information from its neighbouring nodes. For this it is often referred to as message passing. GNNs are able to gather local and global information from a graph. They are able to capture complex relationships like bonding patterns, atomic interactions and spatial configurations that are vital in understanding molecules. [11]

In the protein-ligand interaction interpretation prediction, it is much more important to understand how a ligand (drug molecule) binds itself to a protein (target protein). For this we have to understood that, GNNs are able to capture exquisite chemical & structural relationships of atom molecule like hydrogen bonding, hydrophobic interactions, and electrostatic forces. GNNs are powerful in this field since they can capture these relationships by considering a graph that represent these relationships.

Moreover, some advanced versions of GNNs, such as Graph Convolutional Networks (GCN), Graph Attention Networks (GAN), and Message Passing Neural Networks (MPNN), further increases the model’s capacity by incorporating additional features such as attention and hierarchical feature extraction. These features help to improve the overall accuracy of the model by preoccupied on more significant parts of the molecular structure. [12]

Graph Neural Networks is one of a powerful and biologically relevant framework for modelling in molecular systems. Its ability to effectively capture spatial, relational & structural information in a molecule, it’s a significant reason why it is very suitable for protein-ligand interaction prediction.

### 1.4 Existing Research and Novelty of the Study

The application of GNNs in protein-ligand interaction prediction is not a new task. This problem has been explored in various studies. Some of the existing studies are:

- Graph Convolutional Neural Networks for Predicting Drug-Target Interaction,” *journal of chemical information and modeling*, p. 19, 25 october 2019. [1]
- Protein-Ligand Interaction Graphs: Learning from Ligand-Shaped 3D Interaction Graphs to Improve Binding Affinity Prediction,” *bioRxiv*, p. 21, 07 march 2022. [2]
- A Cascade Graph Convolutional Network for Predicting Protein–Ligand Binding Affinity,” *International journal of Molecular Sciences*, p. 14, 14 April 2021. [13]
- Advanced GNN frameworks mechanisms of multi-graph representations have been introduced recently.
- Explainable Graph Neural Networks (XGNNs) For Protein-Ligand Interaction Interpretation, *International Journal for Research Trends and Innovation*, p. a492, January 2026. [14]

Although various studies have been conducted to improve protein-ligand interaction prediction using GNNs, this problem is not completely solved. Some of the difficulties that make this problem challenging are listed below:

- The lack of interpretability in protein-ligand interaction predictions
- The difficulties in incorporating protein-ligand structural information
- The lack of real-world application systems

Although this project is not a new attempt in this problem domain, it can be considered a contribution to this problem because it is designed to:

- Develop a real-world protein-ligand interaction prediction system using GNNs
- Develop a real-world application User Interface (UI)
- Addressing real-world usability and interpretability challenges

Unlike other existing studies that focus primarily on model development with personal studies and they not provide any usable tools, but this work emphasizes the practical implementation by integrating the GNN model into a user-friendly application which can help to researcher, student and doctor also it can be used by pharma industries to develop drugs by analysing Protein-Ligand Interaction (PLI).

### 1.5 Scop of The Work

Here, i have developed a Graph Neural Network-based system for protein-ligand interaction prediction, and its practical applicability has been hindered, particularly in the fields of drug discovery and bioinformatics. The primary focus of my research involves representing protein and ligand structures at a molecular level as node-edge features, enabling the depiction of intricate relationships and structures in a format compatible with the application of deep learning models.

A comprehensive end-to-end system has been designed, developed, and implemented, incorporating Graph Neural Networks to predict protein-ligand interactions, along with a fully functional and practical system that includes a user-friendly interface for interactive use and related tasks.

This study has also evaluated and addressed issues encountered during implementation and development, including handling biological data, model development challenges, and applying Graph Neural Networks to forecast intricate relationships and structures, such as hydrogen bonding, hydrophobic interactions, and electrostatic forces, model training, and making interactive UI for user.

This study has not evaluated or implemented that how a model such as this can be validated through wet lab-based methods, by including in vitro and in vivo studies, and its suitability for use in this way. This study has bridged the gap between theory and practice by developing a functional and applicable system for predicting protein-ligand interactions, and has assessed and implemented Graph Neural Networks for this purpose.

**Fig 1:**
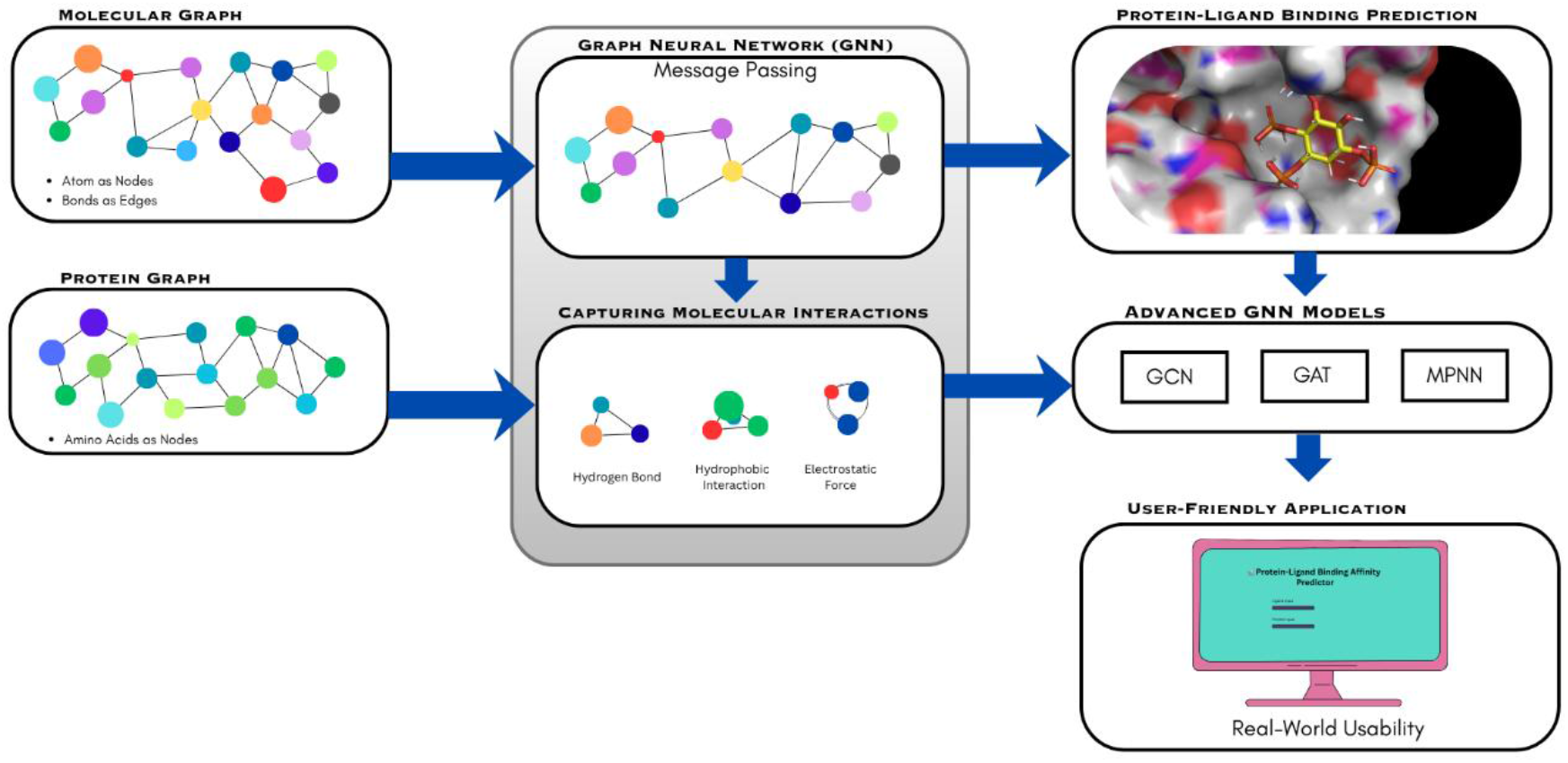
Workflow of the proposed GNN-based protein–ligand interaction prediction system, showing graph construction of molecules, message passing, interaction learning, model prediction using GCN/GAT/MPNN, and integration into a user interface.

## II. Literature Review

### 2.1 Background

Now a days, Protein-ligand interactions (PLIs) prediction plays a central role in modern drug discovery and bioinformatics. A variety of computational methods have been developed to assimilate and forecast these interactions over time. Traditional physics-based methods have revolutionized into advanced Artificial Intelligence (AI)-driven models. Here this part reviews existing literature in a structured way, focusing on key developments, methodologies, and limitations, which provide the basis for the present study.

### 2.2 Traditional Approaches in Protein–Ligand Interaction Prediction

Traditionally, the Protein-ligand interaction prediction acknowledges heavily on traditional computational approaches including molecular docking and molecular dynamics (MD) simulations. These approaches aim to predict the binding orientation and affinity of a ligand with a target protein.

Molecular docking techniques simulate the interaction between a ligand and a protein by finding possible binding conformations and scoring them according to energy functions. Studies such as those by Kitchen et al. in 2004 [15] have shown the effectiveness of docking in virtual screening. Molecular dynamics simulations give insights into the dynamic nature of molecular systems over time, offering a more detailed understanding of binding mechanisms.

These methods have several limitations, despite their importance, such as:

- High computational expense
- Complex systems exhibit limited accuracy.
- The reliance on pre-defined scoring functions.

The challenges have driven the development of data-driven methods.

**Fig 2:**
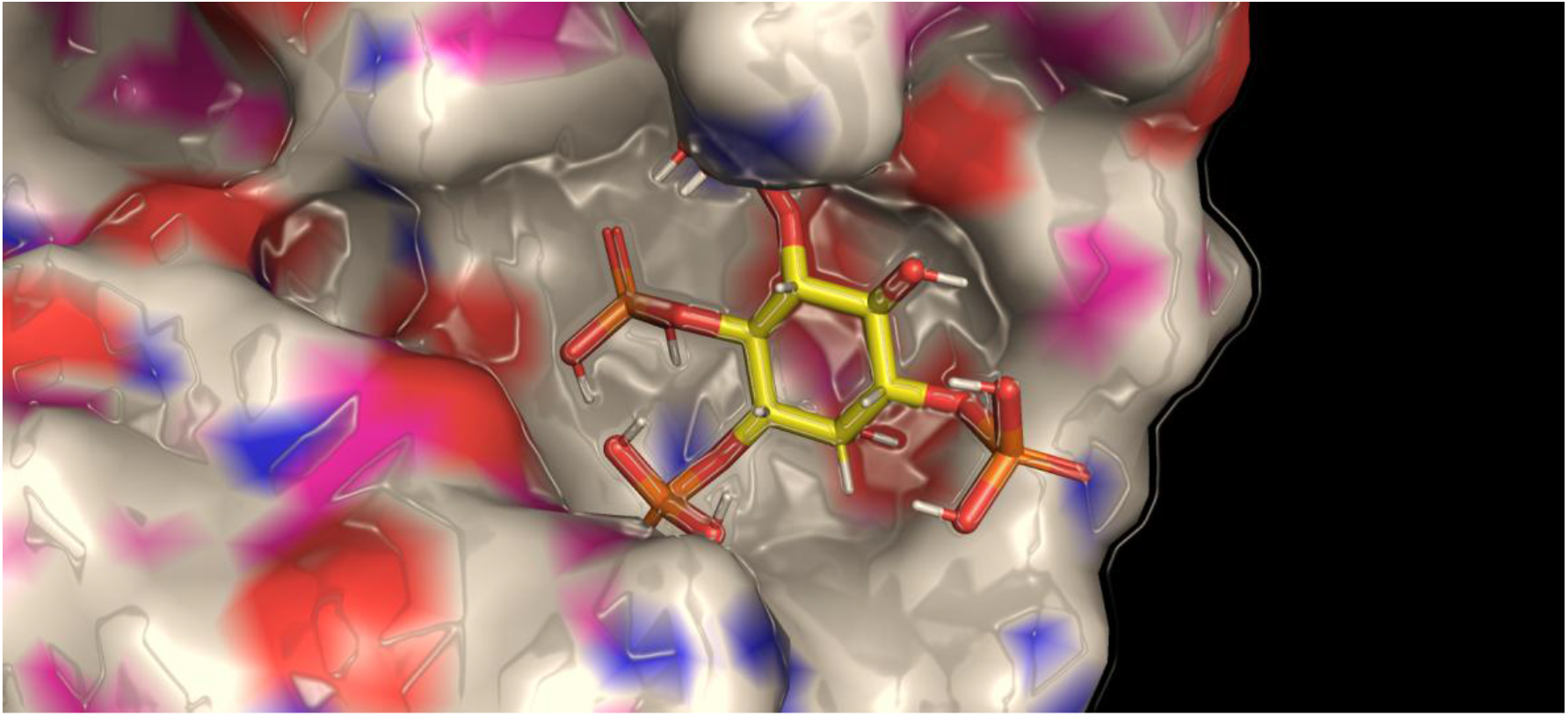
The figure illustrates a traditional approach of Molecular Docking of a Protein with a Ligand molecule.

### 2.3 Machine Learning Approaches in Drug Discovery

The predictive modelling of drug discovery has improved significantly by emergence with Machine Learning (ML). The algorithms of ML like Random Forest (RF) model, Support Vector Machines (SVM) and k-Nearest Neighbours (k-NN) have been used widely to predict the binding affinity & classify drug-target interaction. It can help to reduce the time and cost as compare to traditional methods like Molecular Docking, & MD Simulations.

The ML methods rely some important features extracted from molecular descriptors like:

- Physicochemical properties of the protein
- Molecular fingerprints
- Structural descriptors
- Edge-Node features etc.

Studies like Heikamp and Bajorath (2014) [16] did the application of ML in virtual screening and drug-target prediction.

As like Molecular Dynamics Simulations, ML has some limitations like:

- Dependence on feature engineering
- Inability to capture complex structural relationships
- Limited scalability for large datasets etc.

### 2.4 Deep Learning in Bioinformatics

Bioinformatics has revolutionized by Deep Learning (DL) due to enabling automatic feature extraction from raw data. DL models can learn hierarchical representations directly from input data unlike ML models.

DL models like Convolutional Neural Networks (CNNs) and Recurrent Neural Networks (RNNs) have been applied to protein–ligand interaction prediction. For example, Min, S., Lee, B., & Yoon, S. (2017) [4] used DL in Bioinformatics.

The advantages of DL methods like:

- Automatic feature learning
- Improved predictive accuracy
- Ability to handle large datasets

It faces challenges like:

- Requirement of large training datasets
- Lack of interpretability Difficulty in handling non-Euclidean data structures.

**Fig 3:**
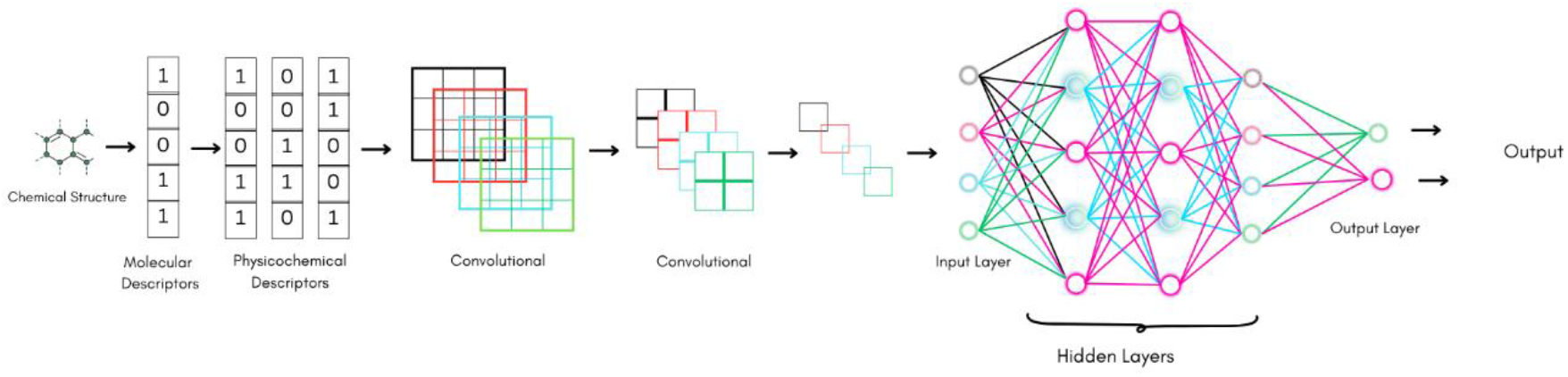
Deep Learning-based pipeline for molecular property prediction, showing conversion of chemical structures into numerical descriptors, feature extraction using convolutional operations, and final prediction through a fully connected neural network architecture.

### 2.5 Graph Neural Networks in Protein–Ligand Interaction Prediction

The Graph Neural Networks (GNNs) are the advanced and emerged solution to handle graph-based data for modelling molecular structures due to their ability. Naturally the molecules can be represented as graph, where atoms are nodes and bonds are edges.

Some studies have demonstrated the effectives of GNNs in drug discovery, such as:

- Wu, Z., et al. (2020) “A comprehensive survey on graph neural networks” [10]
- Zhou, J., et al. (2020) “Graph neural networks: A review of methods and applications” [12]
- A.G.M.G. B. G. Vassilis N. Ioannidis, “GRAPH NEURAL NETWORKS FOR PREDICTING PROTEIN FUNCTIONS,” p. 5, 24 june 2020. [17]
- Duvenaud, D. K., etal. (2015). *Convolutional networks on graphs for learning molecular fingerprints*. NeurIPS. [18]
- Gilmer, J., Schoenholz, S. S., Riley, P. F., Vinyals, O., & Dahl, G. E. (2017). *Neural message passing for quantum chemistry*. Proceedings of ICML. [11]
- Subhasankar Khilar, Dr. Elamathi Natarajan*, (2026). “*Explainable Graph Neural Networks (XGNNs) For Protein-Ligand Interaction Interpretation*” p. a492, January [14]

Recent studies such as Jiang et al. (2021) highlight the ability of GNNs to capture complex molecular interactions & improve prediction accuracy.

## III. Methodology

### 3.1 Overview

This study represents an end-to-end computational framework for predicting protein-ligand interactions from start to finish using Graph Neural Networks (GNNs). The whole methodology used in this work, that combines principles from structural bioinformatics, cheminformatics, and deep learning to model molecular interactions in a graph-based format by using the edge and node (atom and bond) of the compound.

Here, the proposed method is different from traditional approaches like conventional by using data-driven learning to extract structural and relational features directly from molecular graphs, rather than relying on handcrafted descriptors or rigid docking simulations. The workflow comprises several stages, encompassing dataset collection & preparation, graph construction, feature engineering, model design, training, deployment & UI design.

In this work, the GNN model is fully developed and trained by using Python programming language. And all the necessary packages and libraries were installed and imported, majorly like pandas, tqdm, torch, biopython, rdkit, lifelines were used and for result visualization matplotlib is used.

### 3.2 The Foundation of Molecular Representation

In the field of Computational Drug Discovery, the molecules are inherently structured entities, with their chemical behaviour defined by interactions between their constituent atoms. Traditional representations like SMILES strings or molecular descriptors frequently struggle to capture spatial and relational dependencies accurately.

Molecules are represented as graphs to address this:

A graph G = (V, E) where,

- V represents nodes (atom or residues)
- E represents edges (bonds or interactions)

These representation preserves Topological structure, Chemical bonding relationships & Spatial dependencies. Graph-based representations are particularly well-suited for deep learning models designed for non-Euclidean data.

### 3.3 Dataset Acquisition and Preparation

The dataset used in this research work is PDBbind_v2020_refined that consist of experimentally validated protein-ligand complexes obtained from publicly available database that is PDBbind database (PDBbind+ is an enhanced and commercialized version of the original PDBbind-CN website (http://www.pdbbind.org.cn). This upgraded website is a collaborative project between Prof. Renxiao Wang’s team at Fudan University and TopScience Co. Ltd in Shanghai (https://www.tsbiochem.com).

The PDBbind+ web site is officially launched in January 2024. [19]), along with there are more databases are present who consist and provides such kind of dataset to people.

This type of dataset includes 3D structure of protein, protein pocket in pdb format, ligand structure in sdf & mol format. And another very important part is an index file is there that contains the detailed information about all the proteins. That contains information like, PDB code or PDB ID, resolution, release year, −logKd/Ki, Kd/Ki, reference, ligand name of each protein these all information helps to get actual & full dataset.

After getting the dataset, we have to clean & filter it to ensure data quality, and some preprocessing steps are applied such as:

- Removal of incomplete or missing structural data
- Elimination of duplicate entries
- Standardization of ligand representations
- Filtering based on resolution and quality of protein structure.

As the PDBbind dataset do not contains SMILEs code in it, so i generated it by importing Chem from rdkit python library, for that it needs refined file and index file. After many steps of processing i got a csv file that contains all the information of 4620 PDBs & that can help to train the model, that information such as:

i. index_number
ii. protein_pdb
iii. smiles
iv. pkd
v. pki and
vi. p_affinity

**Fig 4:**
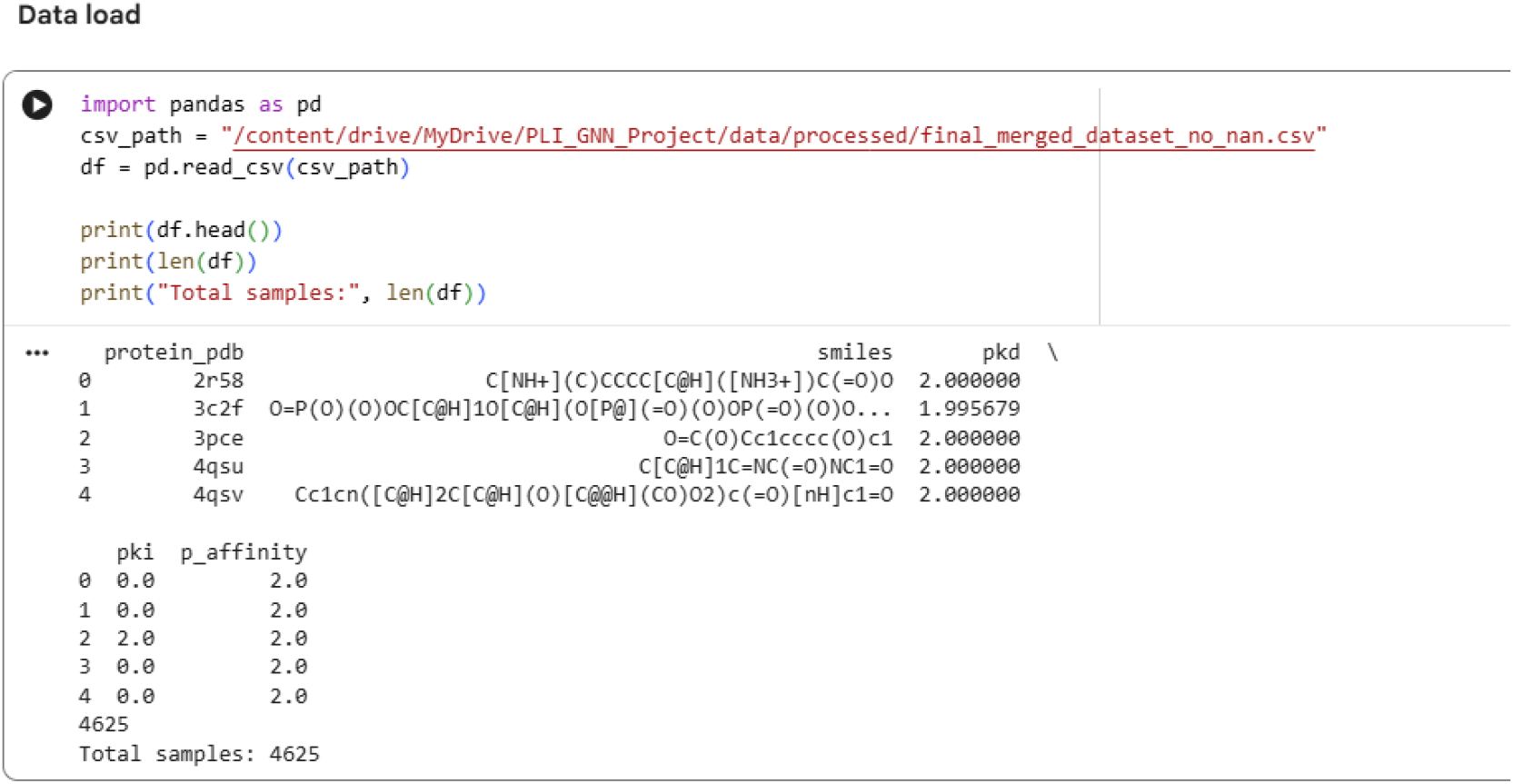
The image shows the organization of dataset after data cleaning

### 3.4 Graph Construction and Encoding

The systematic conversion of molecular structures into graph-based data formats is referred to as graph construction from a molecule, that is G = (V, E), where each node *v* ∈ V corresponds to an atom and each edge **e** ∈ E represents a chemical bond between two atoms. This representation is further enriched with node and edge attributes, including atomic number, hybridization state, bond type, and electronic properties, thereby enabling Graph Neural Networks to effectively learn structural and functional relationships within molecules.

In simply: Atoms → Nodes (Vertices) & Bonds → Edges (Connections).

To construct a graph, we have to consider on Node Features & Edge Features. In Node feature, each atom is encoded with features like: Atom type, Atomic number, Hybridization, Charge, and Aromaticity. Similarly in Edge feature, each bond includes Bond type (single, double, triple), Bond length, Aromatic bond info. And along with one more thing needed that is Graph representation, the final graph G = (V, E, X, A) where,

- V = Nodes (atoms)
- E = Edges (bonds)
- X = Node feature matrix
- A = Adjacency matrix

The next thing is why we need Graph Construction? First thing is the GNN model requires graph input having node and edge features, not raw molecular structures as available. Without graph conversion, the model can’t understand molecule, and the data is unusable for deep learning. Second thing is preserving molecular structure, graphs naturally capture: connectivity, geometry & relationship between atoms. And third thing is instead of manual features like GNN learns patterns automatically detects chemical motifs etc.

#### 3.4.1 Ligand Graph Construction

Ligand molecules were depicted as atom-level graphs, which were obtained from their SMILES notation. In this formulation, each node represents an individual atom and edges symbolise covalent chemical bonds.

Node features included atomic number, aromaticity, formal charge, degree, and valence, thereby capturing fundamental chemical properties along with the whole structure of ligand atoms. The model distinguished between different chemical bonding environments based on edge attributes that encoded bond type, ring membership, and conjugation status. The graph-based representation preserves the molecular topology of ligands, enabling the application of graph neural networks to learn structure– activity relationships pertinent to protein–ligand binding. By using rdkit we ensure that: chemically valid graphs, standardizes feature extraction, reproducibility across datasets this step ensures the model understands chemistry, not just numbers.

#### 3.4.2 Protein Graph Construction

This protein graph construction is simply means that converting to protein’s 3D structures in to a graph representation, where the amino acids or atoms are acting as nodes and their interactions or spatial proximities are represented as edges and its vectors describing their biochemical and structural properties. This feature of Graph Construction enables Graph Neural Networks (GNNs) to capture complex structural & functional relationships among proteins. Another thing is, if the Euclidean distance between their Cα atom was less then 10Å, an undirected edge was created between two residues. The new feature includes a one-hot encoding of amino acid identity, complemented by geometric information derived from residue coordinates.

In my work, 1^st^ I made a list of all amino acids, and after that, i used Kyte-Doolittle hydropathy index and mentioning the charged and polar amino acids by assigning their values.

For Nodes:

- Each node = one amino acid residue
- Only residues from the binding pocket (≤ 8 Å from ligand)

For Edges:

- Distance 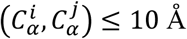

### 3.5 Feature Engineering

Feature engineering involves the systematic or step by step process of extracting and constructing relevant attributes from raw data that accurately capture the underlying patterns necessary for effective predictive modelling. In simple words selection, transformation and creation of relevant input features from raw data to improve the performance and accuracy of Machine Learning model. This process involves techniques like feature selection, transformation, and creation to prepare the input data in a suitable format for machine learning algorithms. In fields like bioinformatics and cheminformatics, feature engineering is essential for encoding biological and chemical data such as molecular properties, structural descriptors, and physicochemical characteristics into numerical formats that models like Graph Neural Networks (GNNs) can process quickly.

The feature engineering transforms raw molecular data into numerical representations that are suitable for model input. It mainly includes Ligand Feature & Protein features. And the Ligand feature have Atom type encoding, Bond type encoding and Molecular fingerprints (Morgan fingerprints). Similarly, the Protein feature includes Amoni acid type (one-hot encoding), Structural properties & 3D coordinates. Mainly the Feature normalization is applied to ensure numerical stability during training.

### 3.6 Graph Neural Network Architecture

The Graph Neural Network (GNN) architecture is a structured design of neural network layers that operates on graph data, in which the information is collected to design the graph by iteratively aggregating and updating features from neighbouring nodes and edges.

Graph Neural Networks (GNNs) is a deep learning framework, specifically designed to process graph-structured data G = (V, E). This model consists of multiple graph convolution or message-passing layers, which iteratively update node representations by aggregating information from neighbouring nodes and edges. Each layer refines the feature embeddings of nodes by capturing both local and global structural patterns. The architecture typically comprises input feature embedding, message passing (aggregation), update functions, pooling/readout layers, and fully connected layers for tasks like classification or regression.

Mainly the core components of GNN architecture are:

a. Input layer
b. Message parsing layer
c. Pooling layer
d. Fully connected layer

a. Input Layer: It is the layer of taking information as input which was already explained previously, that are Node features (X), Edge features (E), and Adjacency matrix (A).
b. Message passing layer: This layer is called as the Heart of GNN model. In which each node collects information from each neighbour, and performs aggregation functions like Mean, Sum, Max. here at each layer K, node representations are updated as :

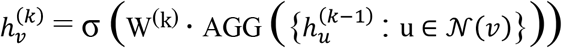 Where:
  - 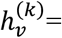 node embedding
  - 𝒩 (𝒱)= neighbors of node *v*
  - AGG = aggregation function (mean/sum/max)
  - σ= activation function
c. The Pooling/Readout Layer: it is also called as Readout Layer, it plays a crucial role in transforming node-level representation, which is required for final prediction task. In each node has its own unique embedding, but prediction is made for the entire graph. So that we need a machine to convert multiple node embeddings to single vector representation. Pooling aggregates these embeddings like:

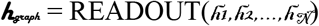
d. Fully Connected Layers: After completion of pooling or readout, the model has a graph-level embedding vector. For the final prediction output, the Fully Connected (FC) Layers are used to map these representations. A Fully Connected layer connects with every input neuron to every output neuron. Like:

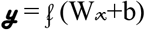

Where:

- *x*= input vector (graph embedding)
- *W*= weight matrix
- *b*= bias
- *f*= activation function (ReLU, Sigmoid, etc.)

### 3.7 Training Strategy

Training strategy refers to the systematic approach or procedure to train a machine learning model, and also includes how the data is prepared, how model learns from it, and hoe the performance is optimized during training. On other hand, training strategy is the complete procedure for optimizing a model’s parameter to achieve accurate predictions. It includes splitting the data, loss function, algorithms of optimization, hyperparameter and protocols of evaluation. In GNN-based protein-ligand interaction prediction model, only the training strategy finalizes that the model effectively earns structural & physiochemical properties from graph representations while avoiding over fittings and increases generalization.

The key components of training strategy:

a. Data Splitting: It simply, division of whole dataset into Training set, Validation set & Test set. In case of Training set, it uses only for learning/training the model, ≈70–80% of data were taken from whole dataset. Here, in this work for training my model, I took 3696 data from whole dataset(4620), that is exactly 80% of my whole dataset, Validation set, it only uses to validate/test the model, that ≈10–15% of data from whole dataset, here I tool 924 data for validate my model that only 20% of data from my whole dataset & Test set, this is only used for evaluate the model, that takes ≈10–15% from whole dataset.
b. Loss Function: Loss function defines how error is measured, for regression (binding affinity) it measures Mean Squared Error (MSE) & for classification it measures Cross-Entropy Loss.
c. Training Process (Epochs & Batching) Training process in GNNs is nothing but it is the iteratively updating model parameters to minimize prediction error, this process is organizing into epochs and batches, enabling efficient and stable learning. An Epoch is the one complete pass or round of entire training dataset through model. During each Epoch, model checks all training sample once. Another main thing is multiple epoch are required for model to learns the patterns effectively of the data. And it requires a neutral number of epoch not too few or not too much. In this work, 150 epochs is used to train the model effectively and each epoch result or value is saved in a separate file.

Batching is unlike epoch, instead of processing the entire dataset at once, the data is divided into smaller subsets. Each batch contains a fixed number of samples (e.g., 32, 64). It reduces memory usage & speed up training. It enables smoother & more stable gradient updates.

## IV. System Development

### 4.1 Introduction

This chapter will describe about design, development of the system for PLI and GNNs. Unlike conventional research that focuses on model development, but this work focuses on the creation of a fully functional, user-friendly application that integrates the trained model into a real-world usable system. Especially this system is designed to make-up the gap between computational drug discovery model and practical usability by permitting users to interact with prediction model through a user-friendly interface.

### 4.2 System Architecture

The proposed system adheres to a modular architecture, a widely accepted design principle in contemporary software and machine learning systems. This architecture breaks down the system into separate yet interconnected modules, facilitating efficient data exchange, scalability, and simplified maintenance. The system consists of three main layers: the frontend (user interface), backend (processing layer), and model layer.

#### 4.2.1 Frontend Layer (User Interface)

This is actually servers as the interaction point among user and system. The only UI responsible for capturing user inputs and giving the prediction results in a clear and interpretable format. In this work the frontend is developed by using Streamlit, which is a lightweight and efficient framework for building data-driven web applications.

The functional responsibilities of the UI is: It accepts the two types of user inputs that are-

- Lead molecule or drug molecule as SMILES code and
- PDB file or PDB ID

User can upload the targeted PDB file otherwise can enter the PDB ID in the given input field.

And next thing is the web application provides interactive controls, like input fields, upload buttons and prediction trigger. After giving all the required data as inputs and hitting the predict button, it will show the predicted Binding Affinity, pKd and pKi value. Along with it will provide a interactive bar plot with showing all the predicted values. Besides it, there are a 3D plot which can be view by rotating with 360^0^.

#### 4.2.2 Backend Layer (Processing Layer)

Every backend layer works as the code processing unit of the whole system. This part is responsible for transforming raw users input into well-structured data suitable for Graph Neural network model. Similarly, as frontend, it also has some functional components:

- Input Validation, it ensures the correctness of uploaded file which is given by the user, checking the valid formats (PDB, SMILES) and handles missing or corrupted inputs.
- Data Preprocessing, this function standardizes molecular structures, cleans & filters input data, & converts raw inputs into consistent formats.
- Graph Construction, this step converts molecules into graph representations, which converts atoms or residues into Nodes and bond or interactions into Edges.
- Feature Extraction, it extracts the relevant features such as, Atomic properties, Chemical descriptors & Structural information. It encodes features into numerical vectors.

**Fig 5:**
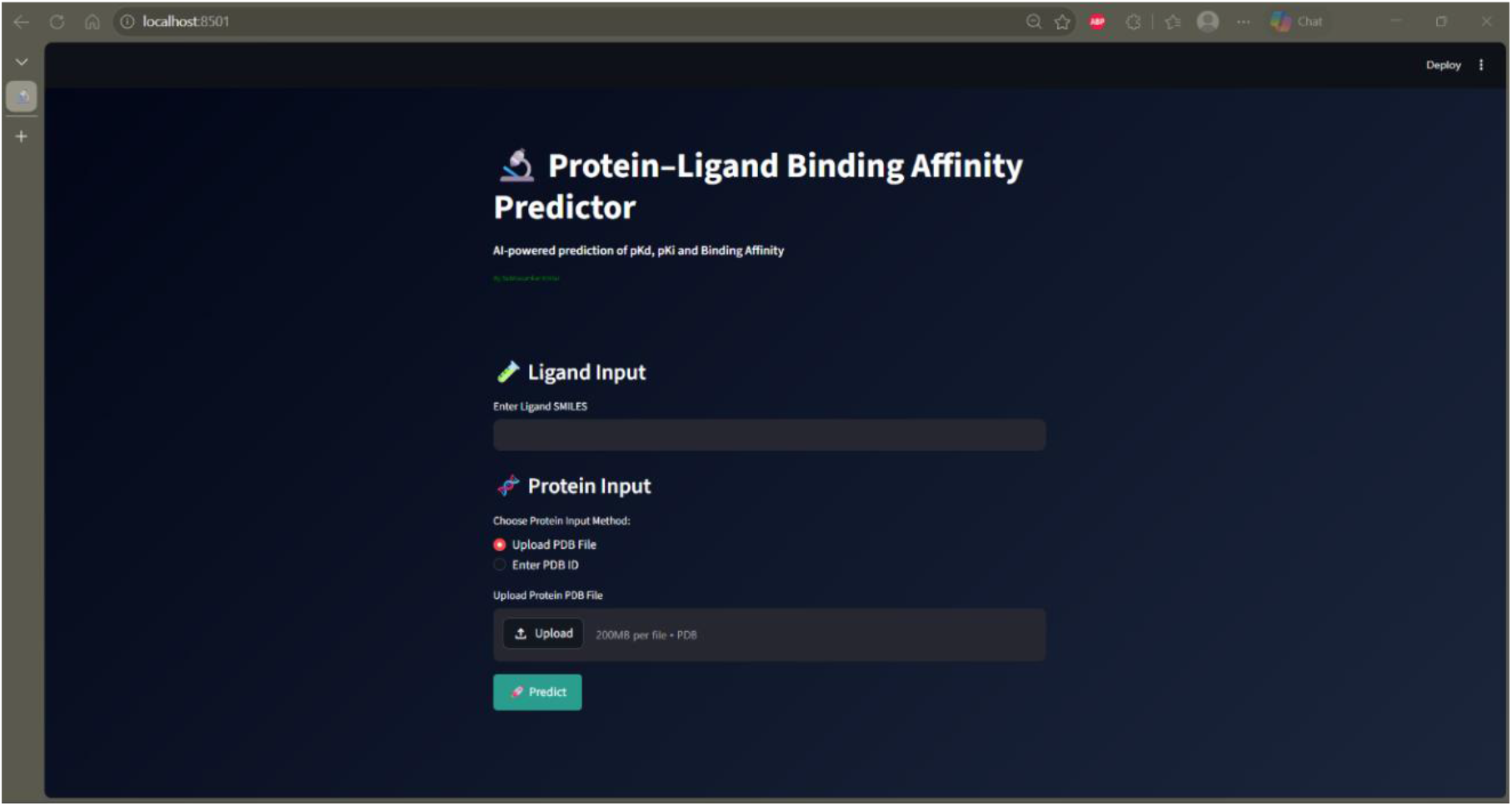
It is the user-friendly UI of developed model, on which user can give Ligand as SMILES code and can give PDB ID or PDB file can be upload

#### 4.2.3 Model Layer (Prediction Layer)

This layer also called as the intelligence core of the system, where the actual prediction is done by a trained Graph Neural Network, which was trained by the real dataset. The trained GNN model is integrated into the application for real-time prediction. This model was developed by using various packages of Python programming language. Here also some core functionalities are there:

- Core Functionalities, it loads the pre-trained GNN models, this layer accepts processed graph data from backend, performs the forward propagation, and generates prediction output.
- Prediction Workflow, the workflow works with forward propagation, step by step Graph Input → GNN Layers → Node Embeddings → Pooling → Fully Connected Layers → Output.
- Model Output, it works on regression output as it produces binding affinity score.

## V. Challenges and Problem-solving

### 5.1 Overview

Development of Graph Neural Network (GNN) based system for PLI prediction involves multiple stages, it starts from data collecting, data handling, model development, and system integration. During the work there are several technical & practical challenges were encountered. Now this section explains these challenges along with the strategies and solutions implemented to overcome them. Along with focusing these aspects illustrate practical complexity of work & contributes a deeper understanding of reals-world system development.

### 5.2 Data Related Challenges

All the biological data especially protein structures often contain different type of noisy & incomplete data in the dataset. That are like: missing residues, incomplete ligand information, low resolution structures etc. to solve this type of dataset problems we have to filter the dataset based on quality criteria, remove incomplete entries and standardize molecular formats.

In the work we need two different types of data, PDB and SMILES this created compatibility issue during processing. So, to solve this issue RDKit was used for ligand processing, converted all inputs into unified graph representations, and built preprocessing pipeline for format handling.

### 5.3 Graph Construction Challenges

There was a major problem occurred in defining edges in protein graph, unlike ligands, proteins don’t have any explicit bond definitions between residues. To overcome this problem distance-based thresholding used, and connected residues in within a specific spatial distance.

Another problem was, handling large graphs, generally the protein structures can be very large, that leading to high memory usage which tends to slow computation. For this problem, I used limited graph size, optimized data structures & efficient batching during training.

### 5.4 Feature Engineering Challenges

Selecting or choosing a relevant feature for atom and amino acid residues was challenging due to the complex physiochemical & structural properties of biological system. So to fix this issue, I selected standard chemical descriptors, used domain knowledge for feature selection such as atomic number, hybridization & aromaticity were considered. Additionally, applied normalization techniques to ensure numerical stability and consistency across input data.

### 5.5 Model Development Challenges

At the time of initial training phase, the model exhibited instability, with loss function failing to decrease consistently. To solve this problem, some optimization strategies were implemented. Learning rate was carefully tuned. Regularization techniques such as dropout were used to prevent over-reliance on specific features.

Another significant challenge was overfitting, where as the model performed well on training data but it showed poor generalization on unseen test data. This issue is normal in Deep Learning models, specially while dealing in complex datasets. To overcome this, the dataset was divided into training, validation, & test set. By doing this the model complexity was reduced by numbers of layers ensuring better generalization capability.

### 5.6 Training Strategy & Epoch Optimization Challenges

One of the major challenges was the epoch optimization, initially the model was trained for only 20 epochs, the result obtained from 20 epoch was not satisfactory, as the model gave poor predictive performance and low accuracy. And this indicated that it had not learned sufficiently the molecular interaction patterns. This is called underfitting where the models fail to capture the complexity of the data.

To address this issue, the number of training epochs were significantly increased 20 epoch to 150 epochs. With extending the train, the model led to more accurate & reliable predictions. However, this improvement came with a computational cost. This 150 epochs took time for 11 hours to train the model from whole data. After 150 epochs were found to be, consistent, stable, and acceptable for practical application.

### 5.7 System Integration Challenges

Integration of trained GNN model with the frontend application possess challenges in maintaining seamless communication between components. To issue is addressed by adopting modular coding approach. Separate functional modules were developed for preprocessing, graph construction and prediction. This ensures smooth interaction between backend processing and frontend display.

This system initially experienced delayed in prediction due to overhead of graph construction and model inference. To address this problem, the preprocessing pipeline was optimized to discard redundant computations. Effective data handling was implemented.

## VI. Results & Evaluation

### 6.1 Introduction

This chapter will describe how the experimental results were obtained from the Graph Neural Network based model for Protein-Ligand Interaction. The model performance is evaluated by using multiple quantitative metrics and visual analysis. This result declares the effectiveness of the method in capturing molecular interaction patterns & generating reliable predictions.

### 6.2 Experimental Setup

The work was done by using the developed GNN model with following configurations:

- Model Type: Graph Neural Network (GNN)
- Framework: PyTorch
- Training Epochs: 150
- Loss Function: Mean Squared Error (MSE)
- Training Time: Approximately 11 hours

### 6.3 Evaluation Metrics

Model evaluation is a demanding step to determine the predictive performance, reliability, & generalization ability of the proposed Graph Neural Networks (GNNs) model. In it multiple evaluation metrics are deployed to comprehensively analyse the model’s effectiveness for both regression and classification.

#### 6.3.1 Root Mean Squared Error (RMSE)

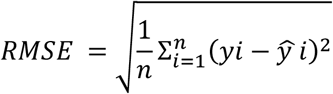

*It measures the square root of the averages squared differences between predicted & actual value. Lower RMSE indicates better model performance and also sensitive to large errors. The RMSE value of my model is 1.424*.

#### 6.3.2 Mean Absolute Error (MAE)

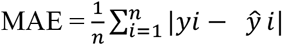

It calculates the average absolute difference between predicted & true value. Unlike the RMSE, it treats all errors equally & less sensitive to outliers. The MAE values is 1.126 has produce from my work.

#### 6.3.3 R^2^ Score

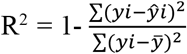

R^2^ shows how well the model explains variance in the data. This function evaluates how the model explains the variance in the target variable. The model produced a value like 0.978 that means it gives nearly perfect prediction.

#### 6.3.4 F1 Score

It is a performance metric used to evaluate classification models by combining precision & recall into a single value. It works as F1 = 2 × Precision * Recall/Precision + Recall. It produced in this work nearly 1, that is 0.9622831787010891.

### 6.4 Evaluation Plots

The Evaluation plots are just graphical tools that used to visualize the data distribution of a dataset. It is easy to compare, analyse & identify the pattern of complex data. Also it help in identification of accuracy, trends & potentials.

#### 6.4.1 Plot 1 (Scatter Plot)

Here the Scatter plot shows True Vs Predicted Binding affinity, the actual (true) pKd values are represented on the x-axis, while the predicted pKd values generated by the model are plotted on the y-axis.

**Fig 6:**
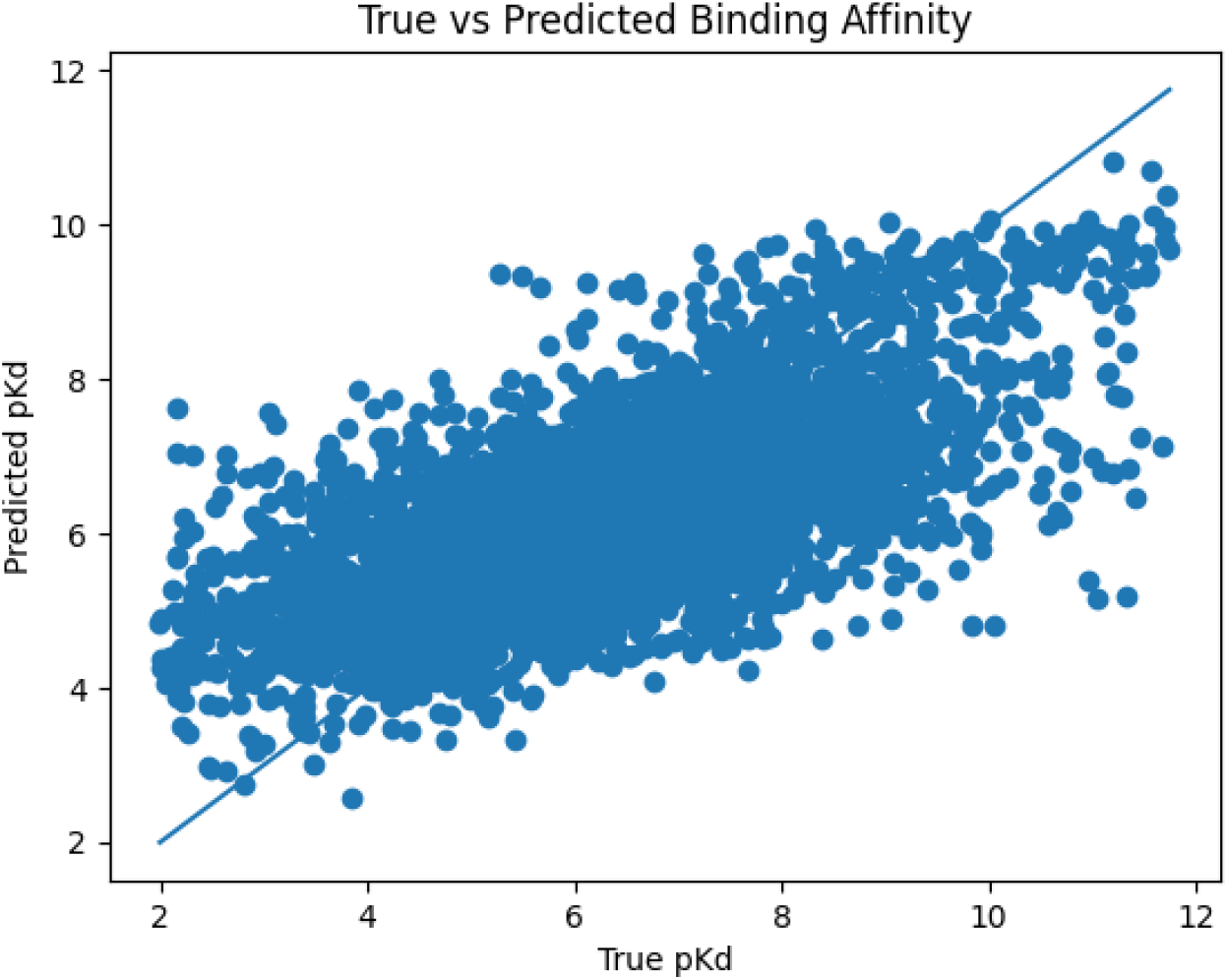
This figure states that, the scatter plot of Binding affinity against Predicted pKd value

##### Analysis

Here the cluster of points are close to diagonal line, it indicates that the model is making accurate predictions. The point distribution shows a positive correlation between true & predicted values, that suggests that the model has successfully learned the relationship between the molecular features & binding affinity. Some dispersion around the diagonal line indicates the presence of prediction errors, which is mostly expected in real world biological data.

#### 6.4.2 Plot 2 (ROC Curve)

The ROC (Receiver Operating Characteristic) Curve is done to distinguishing the positive & negative classes. It plots the True positive rate against False negative rate at different classification thresholds.

Here the blue curve represents the model performance across different thresholds. Whereas the diagonal dashed line represents random guessing or baseline performance. The curve closer to the top-left corner signifies better classification ability. The model archives an AUC of 0.826, it indicates a good discriminative performance. And the AUC value is significantly higher then 0.5, that means the model is much better then random classification.

The results and evaluations were proposed by this GNN-based protein-ligand interaction prediction system. The model performed with low prediction error & stable convergence. The results validate effectiveness of the proposed approach & demonstrate it’s potential for real-world applications in drug discovery.

**Fig 7:**
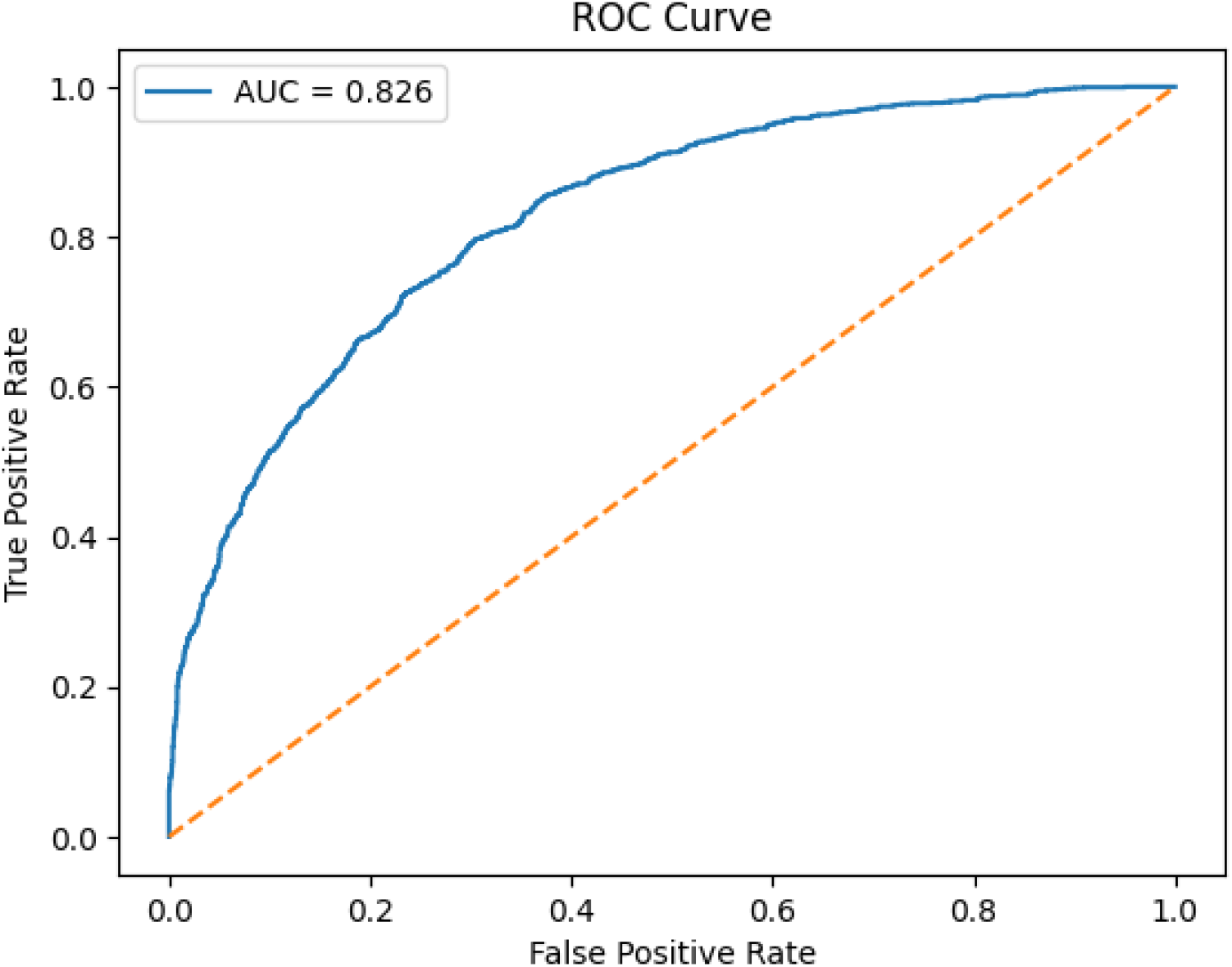
This figure states that, the POC Curve plot of True positive rate against False Positive rate.

## VII. Discussion

This section will provide a depth interpretation of result obtained from proposed GNN-based model base protein-ligand interaction prediction model. Here the discussion focusses on analyse model performance, understanding strength, limitations & comparing the findings with existing literatures. Aim is to calculate the effectiveness of developed system & show its practical importance in drug discovery.

### 7.1 Interpretation of Model Performance

The results & evaluation section demonstrates that, the proposed GNN model is capable of effectively prediction of protein-ligand interactions with satisfactory accuracy. The close difference between predicted & actual value indicates the model successfully learns meaningful representations of molecular structures. The main reason of the GNN model, is to capture Structural & Relational information within molecular graphs. While the traditional models relay on predefined features, whereas the GNN model learns features from the data through message passing mechanisms. This enables the model to understand complex dependencies like atomic interactions & bonding patterns. Furthermore, the improvement of model is determined by increasing the number of Epoh from 20 to 150, it highlights the importance of sufficient training for Deep Learning. Initially, 20 epoch was underfitted, that indicated the model had not yet learned the underlying patterns, so it needed to extend number of epochs to achieve a stable convergence & improve predictive capacity.

### 7.2 Comparison with Existing Methods

Compared with traditional computational approaches like Molecular Docking: Mainly the docking relies on predefined scoring functions. Docking also computationally expensive, time consuming, and limited in capturing dynamic interactions. Here the GNN enters, it learns directly form data, captures complex structural relationships and it can provide faster predictions just after training.

Similarly, compared to conventional machine learning models: the GNN do not require extensive feature engineering, & they preserve molecular structure through graph representation.

### 7.3 Strength of the Proposed System

The proposed system provides several key advantages:

- Graph-Based representation, this preserves molecular structure as node and edges and it captures relational information.
- End-to-End learning, it eliminates need for manual feature engineering.
- Practical Implementation, the system has integrated into a user-friendly application and accessible to non-technical users.
- Scalability, In future, it can be extended by train large dataset, & with advanced models.

### 7.4 Limitations of the Study

- Computational Cost, the model training requires more then 10 hours, if the dataset will be increase, then the training time will increase accordingly. And the resource-intensive process.
- Data dependency, this is one of the main things for model, model performance depends on dataset quality and size.
- Interpretability, the Deep Learning models work like as black-box systems, and it is difficult to explain predictions.
- Lack of experimental validation, we can’t generate the actual experimental data, as it is based on dry lab so all the predictions are computational, and there are no any in vitro or in vivo validations.

### 4.5 Practical Implementations

This developed system has some significant applications for end-user and society, like it can be helpful for Researcher and Scientists, for Drug discovery and health care, for Agriculture and biotechnology etc.

For Scientists and researchers, the model can be integrated with research like Prediction of biding affinity quickly, the don’t need much time or it not consumes more time in the laboratory. It can be use in Virtual screening of candidate molecules, & also it enables rapid identification of promising compounds for further study.

For Drug discovery and Healthcare, this system an be used in Drug development pipeline to prioritize effectiveness of compounds. Personalized medicine, where potential drug responses cab be evaluated computationally. And also, it will support fastest development of treatments for emerging diseases. Another thing is, this leads to reduced drug development time & cost, making medicines more accessible.

For Agriculture & Biotechnology, this developed model can predict interactions between plant proteins & agrochemicals. It can be help to design eco-friendly pesticides & fertilizers. Along with it can help to improve crop protection strategies. This will contribute to sustainable agriculture & food security.

By using this system, we can introduce faster availability of life saving drugs. It can be help to reduce the experimental resources use like chemicals, animals, lab cost etc. can be make a revolution in pharma industry to AI-driven health innovation. It could be making advanced research tools accessible to smaller labs and institutions.

This model is going to be a web-based platform where users can upload their data (SMILES and PDB) and get its prediction. An API service can be integrated with pharmaceutical software pipelines.

## VIII. Conclusion and Future Work

### 8.1 Conclusion

The research work designed and developed for a Graph Neural Network (GNN)-based system for protein–ligand interaction prediction, with a solid description of theory and practical implementation. The main goal was to develop an end-to-end computational work flow having capacity to predict the molecular interactions with a user-friendly interaction and a real-world usability.

The methodology was successfully leveraged graph-based representation of molecules. At where the structure of ligand and proteins are modelled as node and edge. The intention of employing GNN architectures, that can help learn complex interaction patterns. The biological interactions, which includes chemical bonding, spatial relationship, & physicochemical properties, there are essential for understanding the binding mechanism.

The gained results stated that the developed model achieved a stable performance after optimization. At the initial, the model trained for 20 epochs which produced an unaccepted or poor accuracy value, after increasing the epoch value from 20 to 150 the accuracy value improved & the value is accepted. The main or important part of this whole work is integration of the trained model to a user-friendly functional application. Which is developed from using modern tools & frameworks. Unlike other existing studies, that focus on theoretical model development, whereas this project emphasizes practical implementations by integrating webpage an user-friendly UI. This system gives an advanced computational technique for research, student, & professionals in the field.

Overall, this work demonstrated that Graph Neural Networks gives a powerful & effective approach for predicting protein ligand interactions. The combined deployment of both deep learning and real-world application highlights the potential in drug discovery & improves computational bioinformatics workflow.

### 8.2 Future Work

The proposed work gave promising results, and several improvements can be done to enhance the applicability and performance.

- Incorporation of 3D Structural Information: In future work, 3D spatial information of Protein-Ligand complex can be integrated to improve the prediction accuracy and batter capture the molecular interactions.
- Advanced GNN Architectures: The advanced GNN variants such as transformer-based graph networks & attention-based models which are more advanced, they can be used to enhance the model performance.
- Computational Optimization: The system can be more scalable and suitable by reducing the training time and improving the computational efficiency.
- Deployment and Scalability: In future it can be deploy on cloud platforms, enabling access for multiple users & can be ingrate in larger pipeline.
- Developing as a Software: Further works can convert it into a application software which can be help to across all the users.
- Experimental Validation: To validate the model prediction, we can perform in laboratory experiments like in vitro or in vivo, that will give more strength for real-world applicability of the system.

## Notes

https://www.pdbbind-plus.org.cn/download

### Competing Interest Statement

The authors have declared no competing interest.

## References

[1] †. a. R. B. A. W. Torng*, “Graph Convolutional Neural Networks for Predicting Drug-Target Interaction,” journal of chemical information and modeling, no. 25 october, p. 19, 2019.

[2] *. D. K. 2. F. B. 1. C. D. 1. A. B. 3. Marc A.Moesser 1, “Protein-Ligand Interaction Graphs: Learning from Ligand-Shaped 3D Interaction Graphs to Improve Binding Affinity Prediction,” bioRxiv, no. 07 march 22, p. 21, 2022.

[3] M. S. 2. A. K. 3. a. S. D. 4. S. D. 1, “GNNSeq: A Sequence-Based Graph Neural Network for Predicting Protein–Ligand Binding Affinity,” Pharmaceuticals, no. 26 february 2025, pp. 1–31, 2025.

[4] S. L. B. &. Y. S. Min, “Deep learning in bioinformatics,” Briefings in Bioinformatics, vol. 18(5), no. 2017, pp. 851–869, 2017.

[5] J. P. e. a. (. Hughes), “Principles of early drug discovery.,” British Journal of Pharmacology, vol. 162(6), pp. 1239–1249., pp. 1239-1249, 2011.

[6] M. J. Waring, “An analysis of the attrition of drug candidates from four major pharmaceutical companies,” Nature Reviews Drug Discovery, vol. 14(7), pp. 475–486, 2015.

[7] T. &. R. M. Lengauer, “Computational methods for biomolecular docking,” Current Opinion in Structural Biology, vol. 6(3), pp. 402–406, 1996.

[8] Y. B. Y. &. H. G. LeCun, “Deep learning,” Nature, no. 2015, pp. 436–444, 2015.

[9] J. e. a. Vamathevan, “Applications of machine learning in drug discovery and development,” Nature Reviews Drug Discovery, vol. 18(6), pp. 463–477, 2019.

[10] Z. e. a. Wu, “A comprehensive survey on graph neural networks,” IEEE Transactions on Neural Networks and Learning Systems, vol. 32(1), pp. 4–24.

[11] J. e. a. Gilmer, “Neural message passing for quantum chemistry,” Proceedings of ICML, 2017.

[12] J. e. a. Zhou, “Graph neural networks: A review of methods and applications,” AI Open, vol. 1, no. 2020, pp. 57–81, 2020.

[13] Y. Z. 2. C. Z. 3. B. W. 4. a. P. C. 1. H. S. 1, “A Cascade Graph Convolutional Network for Predicting Protein–Ligand Binding Affinity,” International journal of Molecular Sciences, no. 14 April 2021, p. 14, 2021.

[14] Subhasankar Khilar, D. E. N., “Explainable Graph Neural Networks (XGNNs) For Protein-Ligand Interaction Interpretation,” International Journal for Research Trends and Innovation, vol. 11, no. 1, pp. a492–a501, 2026.

[15] D. B. D. H. F. J. R. &. B. J. Kitchen, “Docking and scoring in virtual screening for drug discovery,” Nature Reviews Drug Discovery, vol. 3(11), pp. 935–949, 2004.

[16] K. &. B. J. Heikamp, “Support vector machines for drug discovery,” Expert Opinion on Drug Discovery, vol. 9(1), pp. 93–104, 2014.

[17] A. G. M. G. B. G. V. N. Ioannidis, “GRAPH NEURAL NETWORKS FOR PREDICTING PROTEIN FUNCTIONS,” 2020, no. 24 June 2020, p. 5.

[18] D. K. e. a. Duvenaud, “Convolutional networks on graphs for learning molecular fingerprints,” NeurIPS, 2015.

[19] http://www.pdbbind.org.cn, “PDBbind,” PDBbind, 2024.

[20] J. J. P. Y. F. Hu†, “Interpretable Prediction of Protein-Ligand Interaction by Convolutional Neural Network,” International Conference on Bioinformatics and Biomedicine (BIBM), no. 20 June 2022, p. 4, 2022.

[21] J. S. S. S. R. P. F. V. O. &. D. G. E. Gilmer, “Neural message passing for quantum chemistry,” Proceedings of ICML, 2017.

